# Large particle fluorescence-activated cell sorting enables high quality single cell RNA-sequencing and functional analysis of adult cardiomyocytes

**DOI:** 10.1101/654954

**Authors:** Suraj Kannan, Matthew Miyamoto, Brian Lin, Renjun Zhu, Sean Murphy, David Kass, Peter Andersen, Chulan Kwon

## Abstract

**Rationale:** Single cell RNA sequencing (scRNA-seq) has emerged as a powerful tool to profile the transcriptome at single cell resolution, enabling comprehensive analysis of cellular trajectories and heterogeneity during development and disease. However, the use of scRNA-seq remains limited in cardiac pathology owing to technical difficulties associated with the isolation of single adult cardiomyocytes (CMs).

**Objective:** We investigated the capability of large-particle fluorescence-activated cell sorting (LP-FACS) for isolation of viable single adult CMs.

**Methods and Results:** We found that LP-FACS readily outperforms conventional FACS for isolation of struturally competent CMs, including large CMs. Additionally, LP-FACS enables isolation of fluorescent CMs from mosaic models. Importantly, the sorted CMs allow generation of high-quality scRNA-seq libraries. Unlike CMs isolated via previously utilized fluidic or manual methods, LP-FAC-isolated CMs generate libraries exhibiting normal levels of mitochondrial transcripts. Moreover, LP-FACS isolated CMs remain functionally competent and can be studied for contractile properties.

**Conclusions:** Our study enables high quality dissection of adult CM biology at single-cell resolution.

## INTRODUCTION

The cardiomyocyte (CM) is the primary contractile cell of the heart, and regulation of CM activity is required for coordinated maintenance of proper cardiac pump function. In various pathological conditions, CMs activate initially adaptive cascades of gene changes that eventually lead to cellular dysfunction^1^. These maladaptive genomic changes manifest as heart failure at the organ level and impaired contractility at the CM level. Thus, there is significant interest in elucidating the gene regulatory networks underlying CM remodeling in disease. The advent of single cell transcriptomic methods, particularly single cell RNA-sequencing (scRNA-seq), has offered great opportunities for studying cardiac pathology at single cell resolution^2,3^. This resolution is particularly important as disease processes likely unfold heterogeneously across CMs in the heart^4^. scRNA-seq, in combination with powerful new technologies for rapidly generating genetic mosaic knockouts^5,6^, may enable improved dissection of gene regulatory networks at the single CM level during pathogenic events.

To date, however, scRNA-seq of adult CMs has been limited owing to technical difficulties in the isolation of single CMs. Specialized techniques are required for dissociation of the heart without damaging cells, as CMs are highly sensitive to pH, buffer ionic concentrations, and mechanical disruption^7^. Moreover, adult CMs are large (approximately 125×25µm^8^), rod-shaped cells whose structure precludes isolation by either fluorescence-activated cell sorting (FACS) or commercial single cell microfluidic platforms (such as the Fluidigm C1 or Chromium 10X)^2,3^. Previous studies that have attempted to isolate CMs using conventional FACS equipment have resulted in frequent clogging of the instrument^9^ or generation and/of fragmented myocytes^10^. Moreover, scRNA-seq libraries generated from CMs following FACS^10^ or isolation on the Fluidigm C1^4^ have been dominated by mitochondrial reads, potentially suggesting that the cells were damaged prior to library generation. While one group has previously reported on isolation of rod-shaped CMs by FACS^11,12^, this approach only worked with fixed cells, and required significant instrument and sort parameter customization that may preclude its use in other labs. One approach that has been successful in generating high quality scRNA-seq libraries from single CMs has been single cell picking by micropipette^13^. However, this approach is technically challenging, low throughput, and time intensive, which may in turn lead to significant transcriptomic changes during isolation. In lieu of whole cell scRNA-seq, others have utilized single nuclear RNA-seq (snRNA-seq), which overcomes the challenges of isolating large CMs^14–16^. While this approach can broadly capture cellular heterogeneity, snRNA-seq inherently detects fewer molecules and may skew gene expression towards transcripts predominantly localized in the nucleus (e.g. lncRNAs)^15^. The presence of multinucleated myocytes may confound the analysis as well. Thus, whole cell RNA-seq would be preferable for studying perturbed pathways in disease, particularly where affected transcripts are largely localized in the cytoplasm.

Here, we report on successful isolation of viable single CMs through use of large particle FACS (LP-FACS). LP-FACS is specifically designed for the isolation of large objects, including *C.* elegans, *D.* melanogaster embryos, and large cell clusters. However, use of LP-FACS is not widespread in cell biology applications and particularly with regards to scRNA-seq. We demonstrate that LP-FACS-isolated CMs remained structurally competent and maintained their contractile function. In addition, LP-FACs-sorted CMs generated high quality scRNA-seq libraries. Unlike previous methods, large particle FACS (LP-FACS) is highly user-friendly, and enables rapid, high-throughput isolation of CMs.

## METHODS

Detailed methods, including settings used for sorting, protocols for Langendorff dissociation and scRNA-seq, and methods to produce each figure are included in an online only supplement. We are happy to provide any data, raw or processed, or further information at reader request.

### Experimental Animals

All animal studies were performed in accordance with institutional guidelines and regulations of the Animal Care and Use Committee at Johns Hopkins University. For most experiments, we used adult (>3 months of age) C57L/6J mice from Jackson Laboratory. For mosaic experiments, we used the Ai9(RCL-tdT) mouse from Jackson Laboratory; we refer to this mouse as the Ai9 mouse in the manuscript. To generate mosaics, we performed subcutaneous injection of AAV9-cTNT-EGFP-T2A-iCre-WPRE (Vetor Biolabs) at P1, and analyzed hearts at 8 weeks of age.

### FACS and LP-FACs

We performed conventional FACS experiments using a Sony SH800S cell sorter with both 100 and 130 µm sorting chips. We performed LP-FACS using the Union Biometrica COPAS Flow Platform (FP). The COPAS FP large particle sorter is available in several models, each with a uniquely engineered fluidic path and differently sized flow cell for optimized sorting of different object size ranges. Given that the average CM is 125×25µm, we tested sorting on the COPAS SELECT (now known as the FP-500), which has a flow cell size of 500µm. This ensured that fragile CMs could pass easily through the flow cell regardless of the axis along which the CMs were oriented, and reduced the likelihood of clogs forming in the fluidic path. Detailed settings for both instruments are provided in the supplementary information.

### Langendorff Dissociation

Langendorff dissociation was performed by horizontal cannulation of the aorta and perfusing with a digestion buffer that consisted of Type II Collagenase (Worthington) and Protease (Sigma), as adapted from previous protocols^17–19^. For functional experiments, cells were gradually stepped up to a final Ca^2+^ concentration of 1 mM.

### CM Functional Analysis

CMs were sorted directly onto laminin-coated 7 × 10 mm coverslips and transferred to an inverted microscope (Nikon Eclipse TE-2000U). Cells were loaded with the leak-resistant Ca^2+^ indicator Fura-2AM (Molecular Probes), and analyzed using the IonOptix imaging system and IonWizard software (IonOptix).

### Ethanol Fixation

Ethanol fixation was performed by resuspending pelleted myocytes in 200 uL of cold phosphate buffered saline (PBS), and adding ice cold 100% EtOH drop by drop with gentle stirring to prevent cell clumping.

### scRNA-seq Library Preparation and Analysis

We sorted cells into 96 well plates, which were immediately centrifuged, placed on dry ice, and subsequently stored at −80C. Libraries for scRNA-seq were prepared using the previously established SCRB-seq protocol^20^. We sequenced finished libraries as paired-end reads on an Illumina NextSeq500. Reads were mapped and counted using zUMIs^21^, which uses STAR^22^ two-pass mapping and featureCounts through Rsubread to count. For the Delaughter et al. and Nomura et al. datasets, we mapped reads using the same approach as with data generated in this manuscript. For the Gladka et al. dataset, we were kindly provided with read and UMI counts by the authors. For the Chevalier et al. dataset, we used the publicly available counts data. To identify cell types, we used SingleCellNet^23^ with the Tabula Muris reference^24^. All other packages used in analysis are detailed in the Supplementary Methods section.

### Equivalence Testing Statistics

To compare the equivalence of functional parameters in pre- and post-sorted cells, we used two one-sided t-test (TOST) comparisons as implemented in TOSTER^25^ package. To define our equivalence bounds, we identified the smallest effect size detectable as statistically significant at α = 0.05 by null hypothesis statistical testing for a given sample size. For example, at a sample size n = 14, the smallest effect size that can be statistically significant is 0.78d, where d is Cohen’s d (mean difference divided by pooled standard deviation). Thus, we set the equivalence bounds for TOST at n = 14 to ±0.78d.

## RESULTS

### Recovery of Rod-shaped, Functionally Viable CMs by LP-FACS

To isolate viable CMs, we performed Langendorff perfusion of cannulated adult mouse heart with collagenase and protease as per previously established protocols^17–19^. This resulted in a good yield of CMs as well as other cardiac cell types (Figure 1A). We next tested whether we could isolate viable CMs in two different approaches. In the first method, the Langendorff isolate (including live and dead CMs as well as other non-CM cells) was immediately sorted. In the second method, we performed several sedimentation steps to enrich for CMs while progressively increasing Ca^2+^ concentration to cytosolic levels. In this method, the pre-sort mixture contained only calcium-tolerant CMs, a small proportion of dead CMs, and few, if any, non-CM cells.

**Figure 1.**
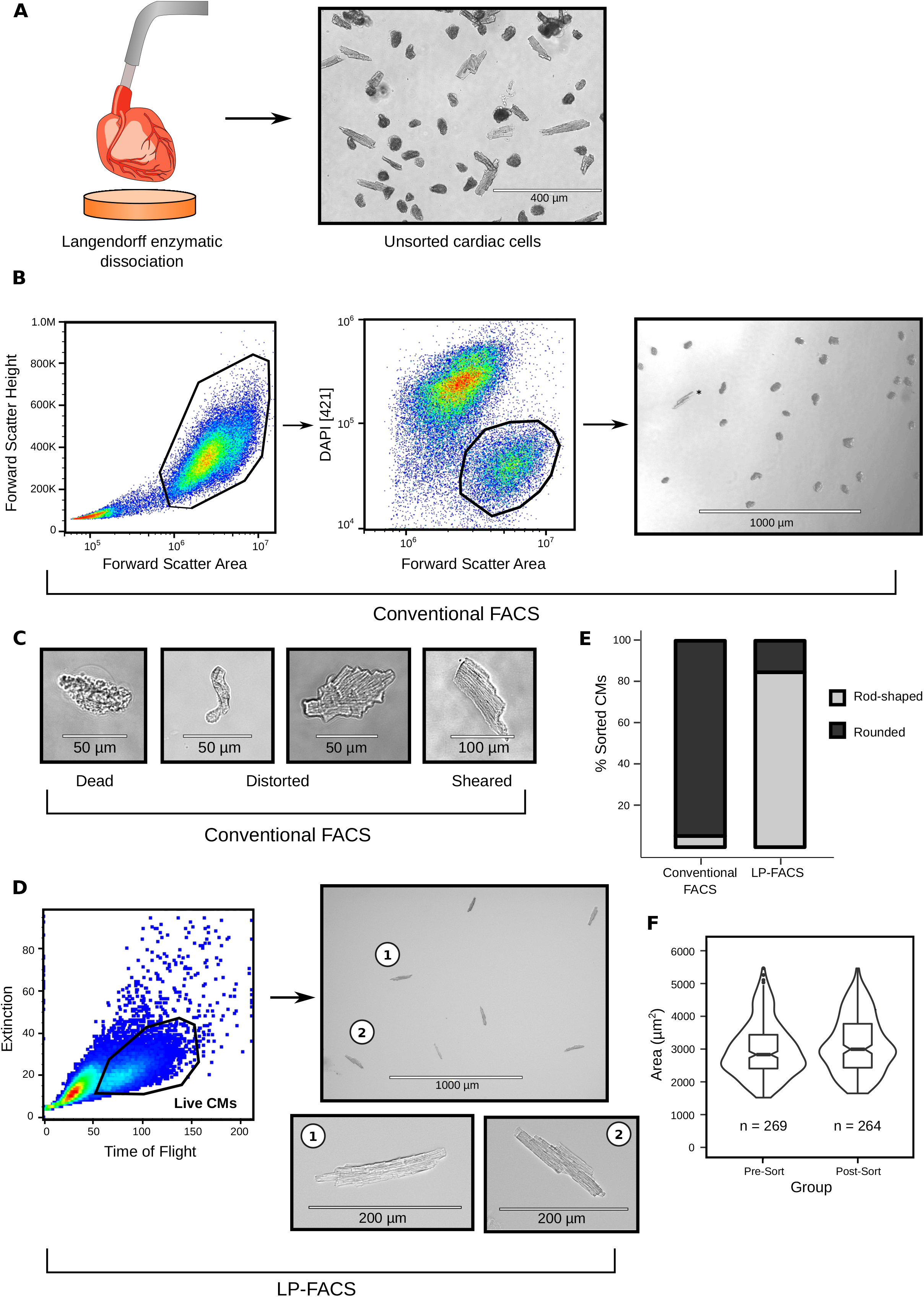
Isolation of CMs from Langendorff-dissociated hearts by conventional and LP-FACs. (A) Retrograde perfusion of collagenase and protease through a Langendorff apparatus produces a mixture of rod-shaped CMs, dead CMs, and other cardiac cell types. (B) Sorting of CMs through conventional FACS. We identified live CMs first through forward scatter and subsequently by selecting DAPI-negative populations. A representative image of this sorted population is shown. The asterisk (*) indicates a rod-shaped CM. (C) Examples of dead or damaged CMs obtained from conventional FACS, representing the vast majority of sorted cells. (D) Sorting of CMs through LP-FACs. We could readily separate live CMs from other cells through time of flight and extinction. A representative image of this sorted population is shown, with higher magnification images of two representative CMs. (E) Percentages of rod-shaped CMs in sorted samples from conventional FACS vs LP-FACS. 200 – 400 cells were analyzed per sorting approach. (F) Comparison of CM area between pre- and post-sorted CMs. Only healthy rod-shaped CMs were analyzed for both groups. N = 3 animals were tested for these experiments.

We first attempted to isolate CMs with conventional FACS, using a commercial cell sorter with standard flow cell sizes (100 µm and 130 µm microfluidic channels). We stained cells prior to sorting with DAPI to readily identify live and dead cells. Despite readily identifying and sorting DAPI-negative CMs, we found that the vast majority of isolated cells were dead, with healthy rod-shaped CMs observed only on rare occasions (Figure 1B). In addition to clearly dead cells, we found numerous cells with sheared or clearly distorted structures (Figure 1C), suggesting that the small flow cell size (relative to CM size) leads to terminal damage of previously healthy CMs.

We next attempted to isolate CMs through LP-FACS. In LP-FACS, a larger flow cell size enables the passage of large objects, and flow settings are optimized to enable gentle isolation of desired objects. As a trade-off, sorting typically happens at a significantly lower rate than conventional FACS (e.g. ten to hundreds of events per second rather than thousands). Here, we performed LP-FACS using a channel size of 500 µm. Through LP-FACS, we found that live, rod-shaped myocytes could be easily separated from other cells with the time-of-flight (measuring axial length) and optical extinction (measuring optical density) parameters (Figure 1D). Notably, we could separate live CMs from dead CMs without the use of a live/dead stain such as DAPI. When sorted, this population was enriched with healthy-appearing myocytes with intact sarcomeric structures. Unlike conventional FACs, where only ~5% of sorted DAPI-negative cells were rod-shaped, we found that ~85% of the LP-FACS-isolated CMs were rod-shaped in appearance (Figure 1E). Moreover, we these cells retained their rod-shaped appearance for several hours post-sort, suggesting that the sorting process did not lead to terminal damage in the sorted CMs. Measurement of CM area pre- and post-sort demonstrates that LP-FACS is capable of isolating CMs across the spectrum of normal size ranges, including relatively larger CMs (Figure 1F). These results support the methodological superiority of LP-FACS for CM sorting.

### Isolation of Fluorescent CMs from a Cardiac Mosaic Model

Conventional gene knockouts in the heart can precipitate cardiac dysfunction and heart failure, producing secondary effects that potentially confound the cell-autonomous effects of gene perturbation. Thus, mosaic cardiac models allow for dissecting primary effects of gene perturbation at the single cell level. While powerful protocols have been developed for rapidly generating mosaic gene knockouts^5,6^, no effective method exists to separate the mosaic populations. We tested whether fluorescent sorting using LP-FACS would enable separation of populations in a simple proof-of-concept mosaic model. We injected P1 Ai9 mice with AAV-cTNT-Cre to generate hearts with a population of tdTomato^+^ CMs (Figure 2A). In these mice, excision of a floxed STOP casette prevents transcription of CAG promoter-driven tdTomato; thus, successful virus infection produces a tdTomato^+^ cell. We subsequently isolated the hearts at 8 weeks of age and performed Langendorff dissociation. LP-FACS readily separated healthy RFP^+^ and RFP^−^ CMs from one another (Figure 2B), suggesting the utility of the instrument for purification of different cell populations in cardiac mosaic models.

**Figure 2.**
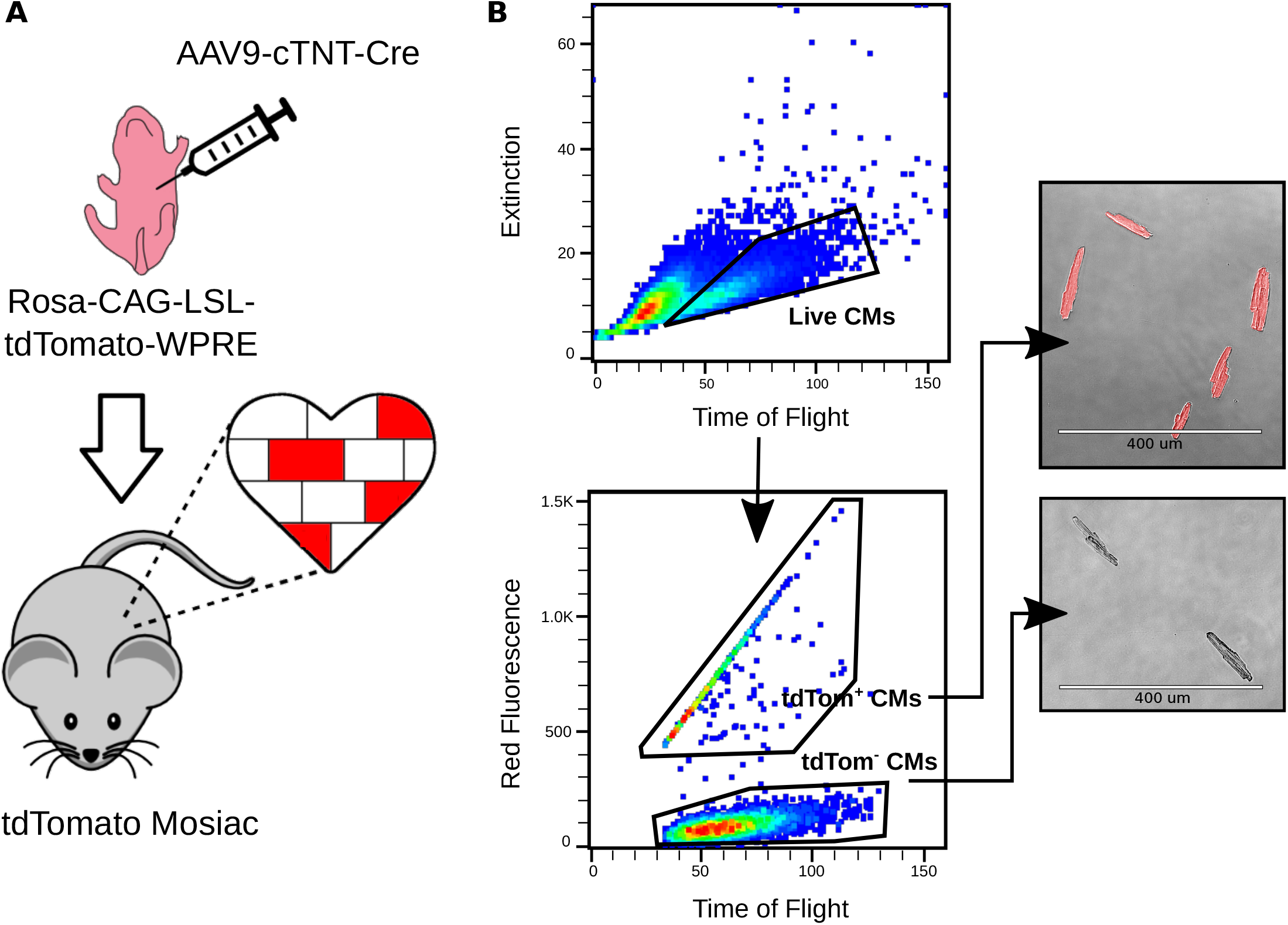
Sorting of CMs from tdTomato mosaic mouse. (A) Workflow for experimental strategy. Animals were analyzed at 8 weeks of age. (B) Gating strategy for sorting tdTomato mosaic hearts, with representative images for each relevant gate. Cells were first filtered for live CMs and subsequently sorted based on red fluorescence.

### Generation of High Quality scRNA-seq Libraries from Sorted CMs

Having validated the successful recovery of structurally competent adult CMs through LP-FACS, we next tested whether we could use isolated CMs for transcriptomic assays. We first isolated RNA from 500, 1000, and 1500 cells respectively, and assessed RNA quality using the Advanced Analytical Fragment Analyzer system. The Fragment Analyser system uses a quantification of the 18S and 28S ribosomal subunit peaks to compute an RNA Quality Number (RQN) that denotes the integrity of input RNA. We found that all three samples presented with 28S/18S ratios between 1.8 and 2.4; correspondingly, RQN was 10.0 for all samples, indicating maximal RNA quality (Figure 3A).

**Figure 3.**
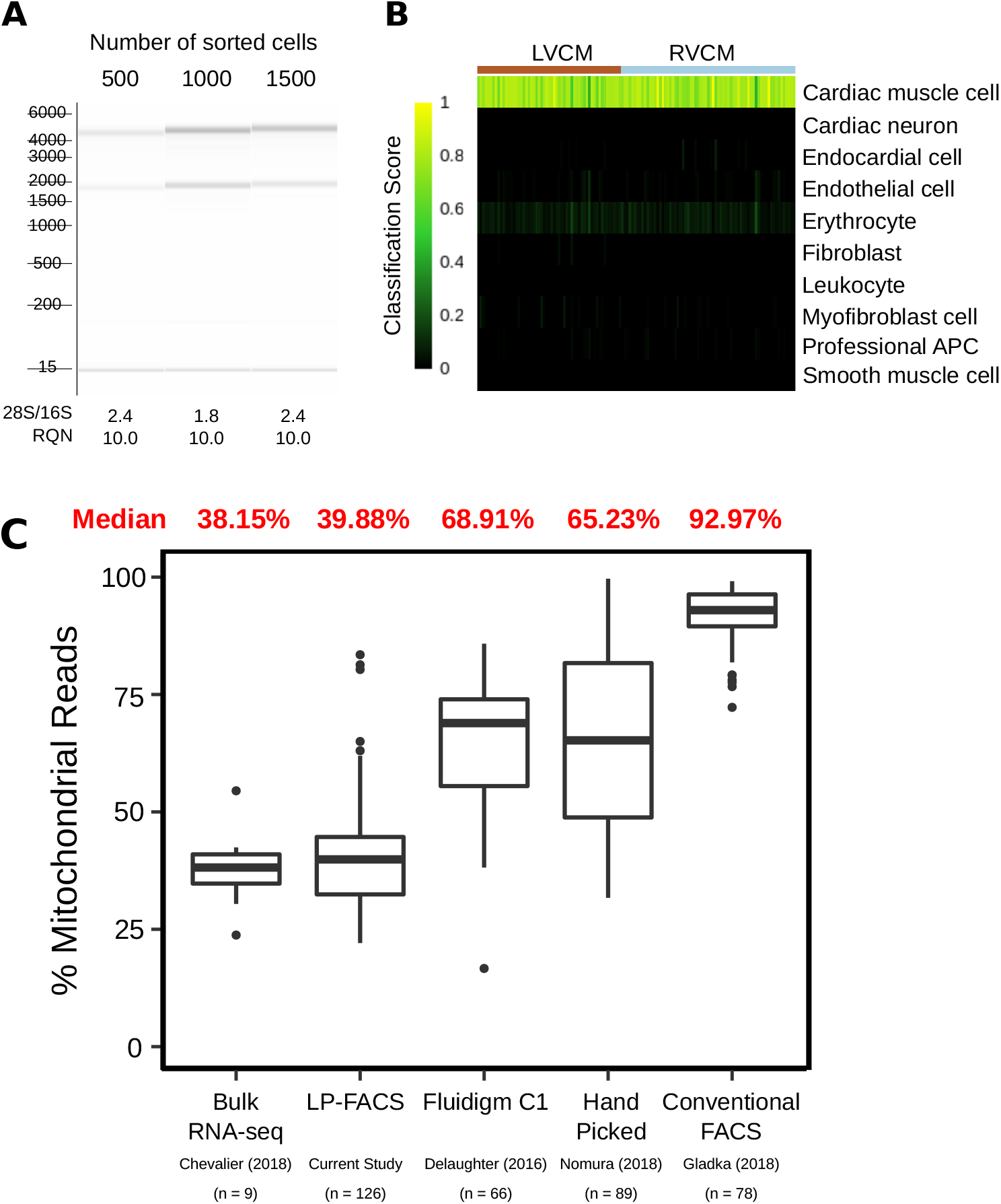
scRNA-seq of LP-FACS isolated CMs. (A) Bioanalyzer gels for 500, 1000, and 1500 sorted CMs. 28S/16S and RQN values are provided below. (B) Output of SingleCellNet classification of isolated CMs. LVCM = left ventricular CM, RVCM = right ventricular CM. (C) Comparison of percentage of mitochondrial reads in LP-FACS-isolated CMs vs. one bulk study and three other single cell studies using different isolation methods.

Encouraged by these results, we performed a proof-of-concept scRNA-seq experiment on LP-FACS-isolated adult left ventricular (LV) and right ventricular (RV) myocytes (57 LVCM, 69 RVCM after RNA-seq quality control) using the established SCRB-seq protocol^20^. To validate that our sorted cells were indeed CMs, we used the SingleCellNet tool, which classifies cell types in scRNA-seq datasets by comparison to established cell atlases^23^. We found that all cells were identified with high confidence as cardiac muscle cells (Figure 3B). Indeed, sorted cells showed high expression of known mature CM markers such as *Actc1*, *Tnnt2*, *Tnni1*, and *Myh6* but low expression of markers associated with other cardiac cell types such as endothelial cells, fibroblasts, and smooth muscle cells, or potentially contaminating cells such as erythrocytes (Supplemental Figure 1A).

We next sought to assess the quality of our sequenced CM libraries. One commonly used metric for assessing scRNA-seq quality is the percentage of reads originating from mitochondrial transcripts^26,27^. In addition to being a potential read out of cellular stress, a high mitochondrial read percentage is a useful indicator of potential membrane rupture and cell death. In a damaged cell, cytoplasmic content (and thus cytoplasmic RNA reads) will leak out of the cell, while RNAs in the mitochondria are retained. Thus, a high percentage of reads being mapped to mitochondrial genes indicates likely cell damage. Based on this concept, we compared the percentage of reads coming from mitochondrial transcripts in our sequencing study as well as three other scRNA-seq studies of CMs: DeLaughter et al. (2016)^4^, in which the Fluidigm C1 system was used to isolate p21 LVCMs; Nomura et al. (2018)^13^, in which single cell picking was used to isolate adult CMs; and Gladka et al. (2018)^10^, in which conventional FACS was used to sort adult CMs. As a control, we also compared against Chevalier et al. (2018)^28^, who generated bulk RNA-seq datasets of isolated ventricular CMs. CMs are a highly metabolically active cell type with a high volume of mitochondria; previous studies have indicated that mitochondrial transcripts comprise approximately 30% of total mRNA in the heart^29^, and between 30-50% of transcripts in CMs in particular^28,30^. We found that in our study, mitochondrial transcripts accounted for ~40% of reads, corresponding well with bulk studies (Figure 3C). However, in other studies, mitochondrial transcripts accounted for a notably higher percentage of sequenced reads. These higher percentages indicate likely cellular damage and/or rupture due to harsher methods of CM handling, supporting the use of LP-FACS to generate improved quality CM scRNA-seq libraries. Because CMs are generally mitochondria-rich, concerns have been raised about the ability of scRNA-seq to detect genomic genes in CMs^2^. In our sequencing study, we found that we were able to detect ~3700 genes at a subsampled read depth of 200,000 reads, which is comparable to gene detection of non-CM cell types using the SCRB-seq protocol^31^ and to other current CM scRNA-seq datasets (Supplemental Figure 1B).

### Sorting of Ethanol-Fixed CMs

In certain experiments, it may be infeasible to immediately sort cells immediately after isolation. Thus, we investigated whether we could successfully sort and perform scRNA-seq on ethanol-fixed CMs. We fixed post-Langendorff isolate using 80% ice cold ethanol and found that we were able to recover rod-shaped CMs through LP-FACS while using the same time-of-flight and optical extinction parameters as with live cells (Figure 4A). We then performed scRNA-seq on 36 live and 36 fixed CMs. When downsampling the data to lower read depths (<250,000 reads/cell), we found that live and fixed CMs were similar in terms of detected genes and detected unique molecular identifiers (UMIs), corresponding to total detected molecules in each cell. However, at higher read depths, live cells began to outperform fixed cells both in terms of gene and UMI detection (Figure 4B, C). Even at high downsampled read depths, however, expression of genes in live and fixed cells was highly concordant (Figure 4D). These results suggest that while individual mRNAs may be lost with ethanol fixation, overall gene expression patterns are maintained. In particular, while live cells may be superior for high depth sequencing experiments, fixed cells may be sufficient for large-scale experiments where per-cell read depth is lower.

**Figure 4.**
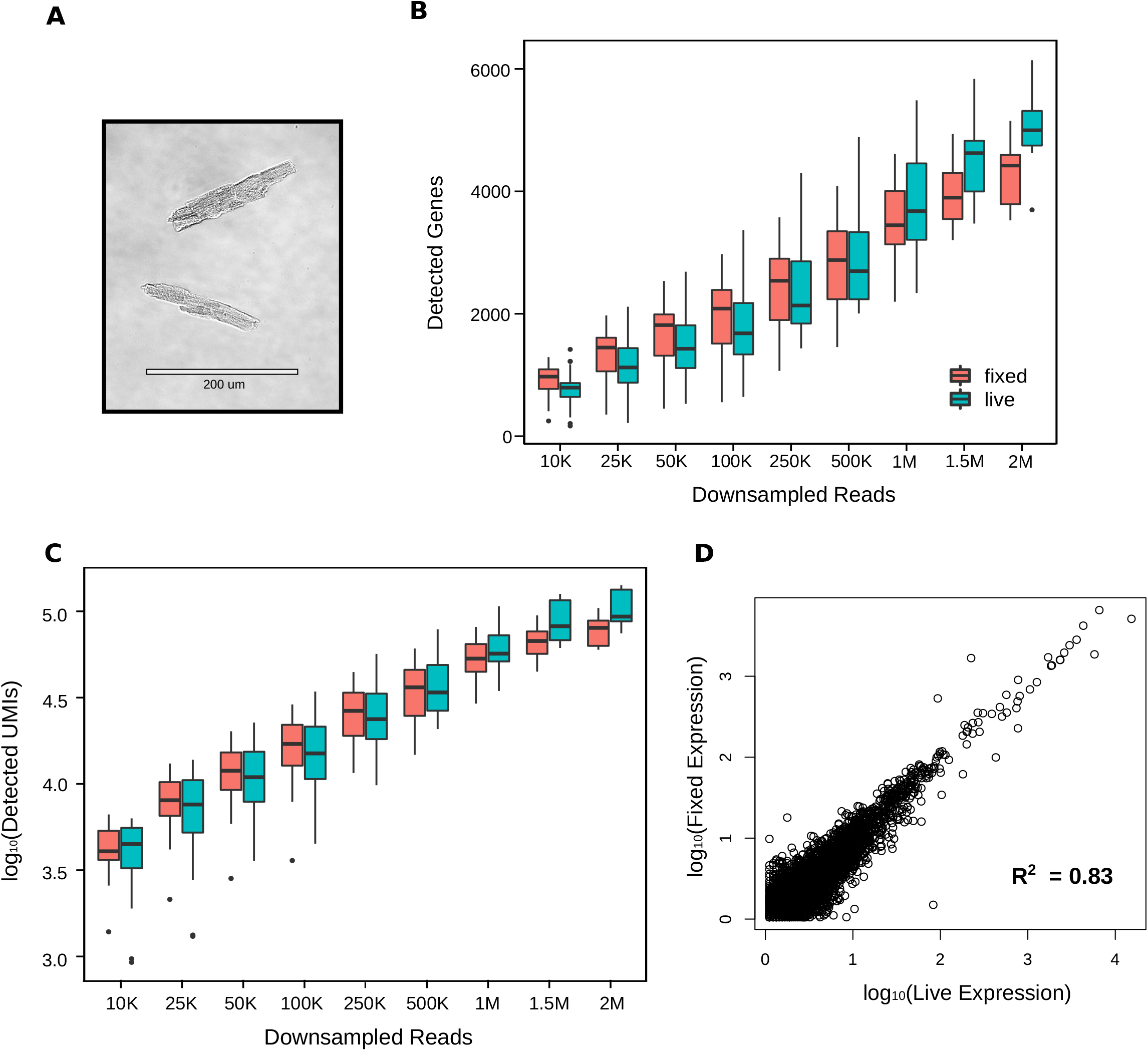
scRNA-seq of LP-FACS isolated fixed CMs. (A) Representative image of fixed CMs post-sort. (B) Number of detected genes at various sampling depths for both live and fixed cells. (C) Number of detected molecules (estimated by unique molecular identifiers, UMIs) at various sampling depths for both live and fixed cells. (D) Correlation of expression levels (in UMIs) between expressed genes in fixed and live CMs at a subsampled depth of 1.5M reads per cell.

### Contractility and Calcium Transient Analysis from Sorted CMs

We next investigated the functional properties of LP-FACS-isolated CMs. We found that calcium-tolerant CMs could be paced post-sort, and maintained sarcomeric shortening and calcium transient traces comparable to pre-sorted CMs (Figure 5A). We further analyzed several parameters of contraction and relaxation (Figure 5B-E) as frequently studied read-outs of CM function. To assess whether pre- and post-sort parameters were equivalent, we utilized TOST, which helps determine whether the difference between two populations falls within some acceptable boundary that supports functional equivalence. We defined a smallest effect size of interest based on the smallest possible effect size detectable as statistically significant (see Methods; raw bounds and calculations are shown in Figure S2). Within these bounds, TOST supported statistical equivalence between pre- and post-sorted cells for several important parameters, including fractional shortening (Figure 5B), change in calcium transient (Figure 5D), and calcium transient decay τ (Figure 5E). However, we did note a difference in relaxation dynamics, particularly as demonstrated by the time to return to 50% baseline during sarcomere shortening (Figure 5C). Nevertheless, the data broadly support the maintenance of contractile function in LP-FACS-isolated CMs, particularly with regards to fractional shortening and calcium transient parameters.

**Figure 5.**
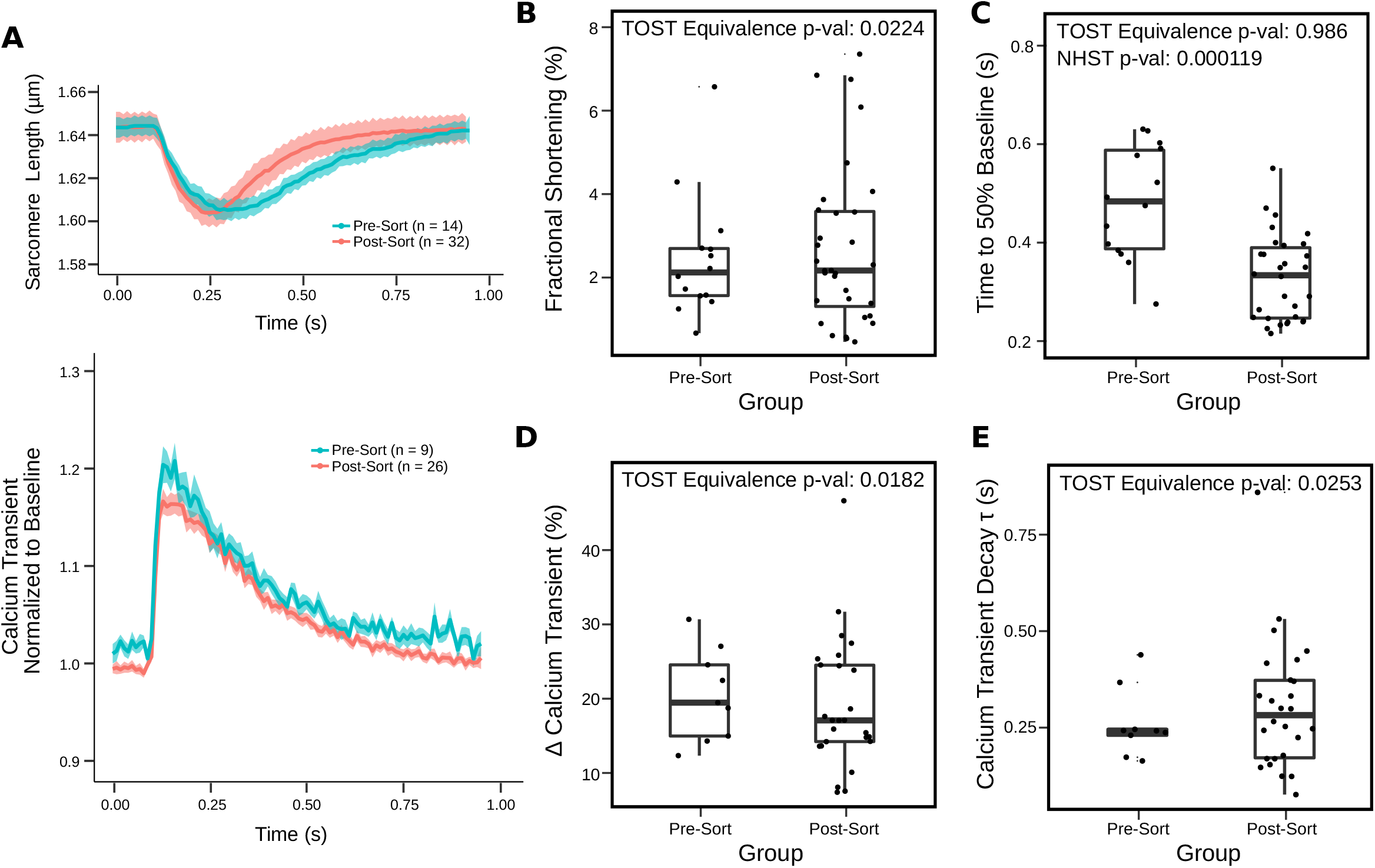
Functional analysis of LP-FACS-isolated CMs. (A) Sarcomere shortening (top) and calcium transient (bottom) traces for pre- and post-sorted CMs. Traces are shown with a thick line representing the mean and shaded are representing standard error of the mean. (B – C) Parameters for sarcomeric shortening. (D-E) Parameters for calcium transients. All measurements were taken in N = 4-6 animals.

## DISCUSSION

Here, we present a method to isolate single CMs through use of LP-FACS. Previous approaches have been limited due to the large length of adult CMs (~125um on average), thus typically yielding damaged or sheared CMs^3^. In line with this, when attempting to sort adult CMs on a commercial cell sorter, we were unable to recover intact CMs and experienced several clogs even when using the largest available channel size (130um). While some groups have established CM sorting protocols using conventional sorters^11,12^, these approaches have required careful calibration of sort and machine parameters, which may not be possible or easily achievable on most types of commercial sorters. By contrast, the COPAS FP system readily isolates healthy and intact adult CMs through LP-FACS with no additional customization. Viable myocytes could be separated from dead cells and non-CMs solely through time-of-flight and optical extinction parameters. While live sorting is preferable for many applications, including high depth scRNA-seq as well as studies requiring downstream single cell functional analysis, LP-FACS also readily worked with fixed CMs. Intriguingly, post-sorted CMs remain functionally competent and can be studied using standard single CM sarcomeric shortening and calcium transient assays. To our knowledge, LP-FACS is the only approach that enables both generation of high-quality scRNA-seq libraries *and* allows for isolation of cells for functional analysis. We envision this technology will enable researchers to integrate transcriptomic and functional data for a range of cardiac disease models, including novel mosaic knockdown models.

We performed scRNA-seq on sorted CMs and found that the sequenced libraries faithfully recapitulated known CM gene expression patterns. A previous concern regarding scRNA-seq of CMs has been the large number of reads mapped to mitochondrial genes, which has approached 90% in some studies^2^ and likely indicates cell rupture. Here, we found that the percentage of reads mapped to mitochondrial genes (~40%) matched bulk RNA-seq datasets of CMs and was notably lower than other scRNA-seq studies, indicating improved library quality. Additionally, despite a relatively high percentage of reads (in comparison to other tissues) being mapped to mitochondrial reads in CMs, we were able to detect genes with similar sensitivity to other cell types using the same sequencing method (SCRB-seq). Nevertheless, fturther reduction in the number of sequenced mitochondrial reads could improve the quality of the LP-FACs isolation approach. Several groups have developed CRISPR/Cas9 based approaches for targeted cleavage and removal of reverse-transcribed mitochondrial RNAs^32,33^. These methods may allow for improved deep sequencing of sorted CMs in future experiments.

Our study represents a significant improvement in the generation of high quality scRNA-seq libraries for adult CMs. However, our proof-of-concept experiments in this manuscript used relatively few cells per experiment. We note that this was not a limitation of the technology but rather to assess the feasibility of our pilot experiments. While large particle sorting inherently requires lower flow rates than conventional sorting, we found that we could acquire several thousand healthy myocytes within 5 – 10 minutes of sorting. Thus, in combination with powerful new split-pool barcoding techniques^34^, LP-FACS could enable generation of large scale CM scRNA-seq libraries.

It is important to observe that successful use of LP-FACS requires a protocol for successfully dissociating cardiac tissues into a single cell suspension that can be sorted. Here, we utilized the well-established Langendorff perfusion system for digesting mouse hearts^35^. This approach has been applied to a variety of small mammal models, and enables reproducible and efficient adult cardiac dissociation, though successful non-Langendorff methods have also been described^7^. For larger tissues such as human surgical biopsy tissue, both chunk-based^36^ and perfusion-based methods^37^ have been applied to successfully yield rod-shaped CMs. We anticipate that future improvements in dissociation methods, particularly for human tissues, will enable improved recovery and quality of single CMs via LP-FACS.

While our initial goal in exploring large particle sorting was to improve scRNA-seq library quality for adult CMs, we were struck by the observation that post-sorted CMs appeared to maintain intact sarcomeric structure when sorted into a non-lysis buffer. We were thus curious to know if sorted CMs could be maintained for functional assays. Our results demonstrate that the large particle sorting process maintained normal CM contractile properties and EC-coupling mechanics. This advantage of using LP-FACS will allow for the study of how broach shifts in the transcriptome affects individual CM function.

One current limitation of the approach detailed in this manuscript is the exclusion of non-CM cell types. The heart is composed of a heterogenous mixture of cell types, each of which contributes to the full range of physiological function of the adult heart. Our focus on CMs was due to the abundance of CMs in the heart, their important biological role, and the failure of other methods to isolate single CMs for transcriptomic analysis. By contrast, previous studies have successfully utilized droplet-based approaches to sequence non-CM cells in the heart^38^. Theoretically, the LP-FACS approach can isolate non-CMs so long as they can be distinguished from dead CMs through other parameters. For example, use of a transgenic fluorescent line to identify CMs, or a fluorescent stain to identify dead cells, could enable simultaneous isolation of live CMs and non-CMs. Additionally, while nearly all CM functional parameters were identical, differences in CM relaxation kinetics were detected. We suspect minor technical differences in protocol contributed to this, and further optimization of sorting conditions, such as applying strict selection criteria of pre-sorted CMs, will eliminate all functional discrepancies. This will be important for studies that seek to determine genetic and functional linkages in heart disease, and studies that seek to establish this are underway. With these optimizations, we believe the approach detailed in this manuscript will assist in the isolation of cardiac cells and enable improved elucidation of gene regulatory networks in the heart in health and disease.

## AUTHOR CONTRIBUTIONS

S.K., M.M., and B.L. were directly responsible for performing experiments in this manuscript. R.Z., D.K., P.A., S.M., and C.K. provided significant intellectual contribution to the manuscript. The manuscript was initially written by S.K. with subsequent revisions and input from all authors.

## ACKNOWLEDGEMENTS

We thank Dr. Deborah Andrew for kindly allowing us to use her lab’s COPAS instrument. We additionally thank Kevin Mangs for technical support regarding the COPAS instrument, and Julia Thompson and Mike Fazzio for answering questions regarding scientific applications of the COPAS FP. We additionally thank Grace K Mueller for helpful feedback on several experiments. We lastly thank Bas Molenaar and Dr. Eva von Rooij for kindly providing us with the data from their single cell study. This work was supported by grants from NICHD/NIH, AHA, and MSCRF.

---

**Figure S1A.**
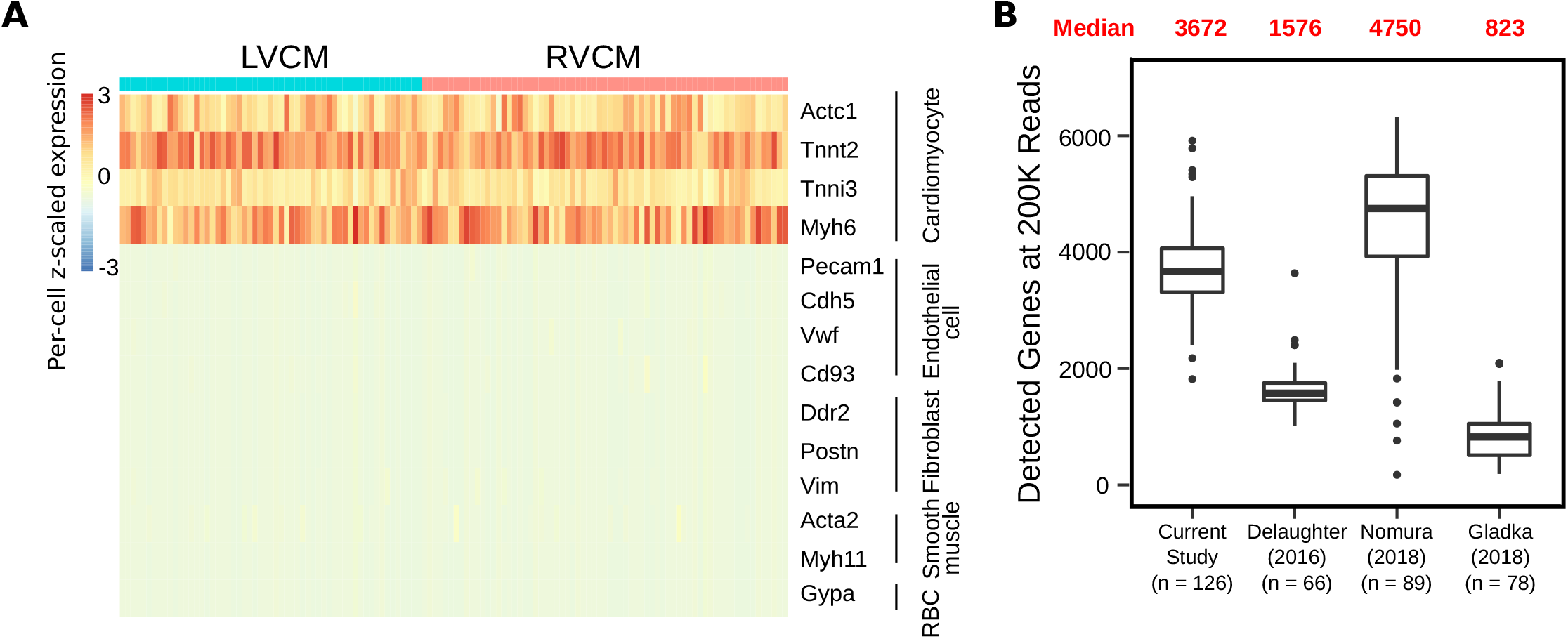
scRNA-seq of LP-FACS isolated CMs. (A) Heatmap of candidate genes representing important cardiac cell types. LVCM = left ventricular CM, RVCM = right ventricular CM. (B) Comparison of detected genes in LP-FACs-isolated CMs vs CMs from other studies at a subsampled depth of 200K reads per cell.

**Figure S2A.**
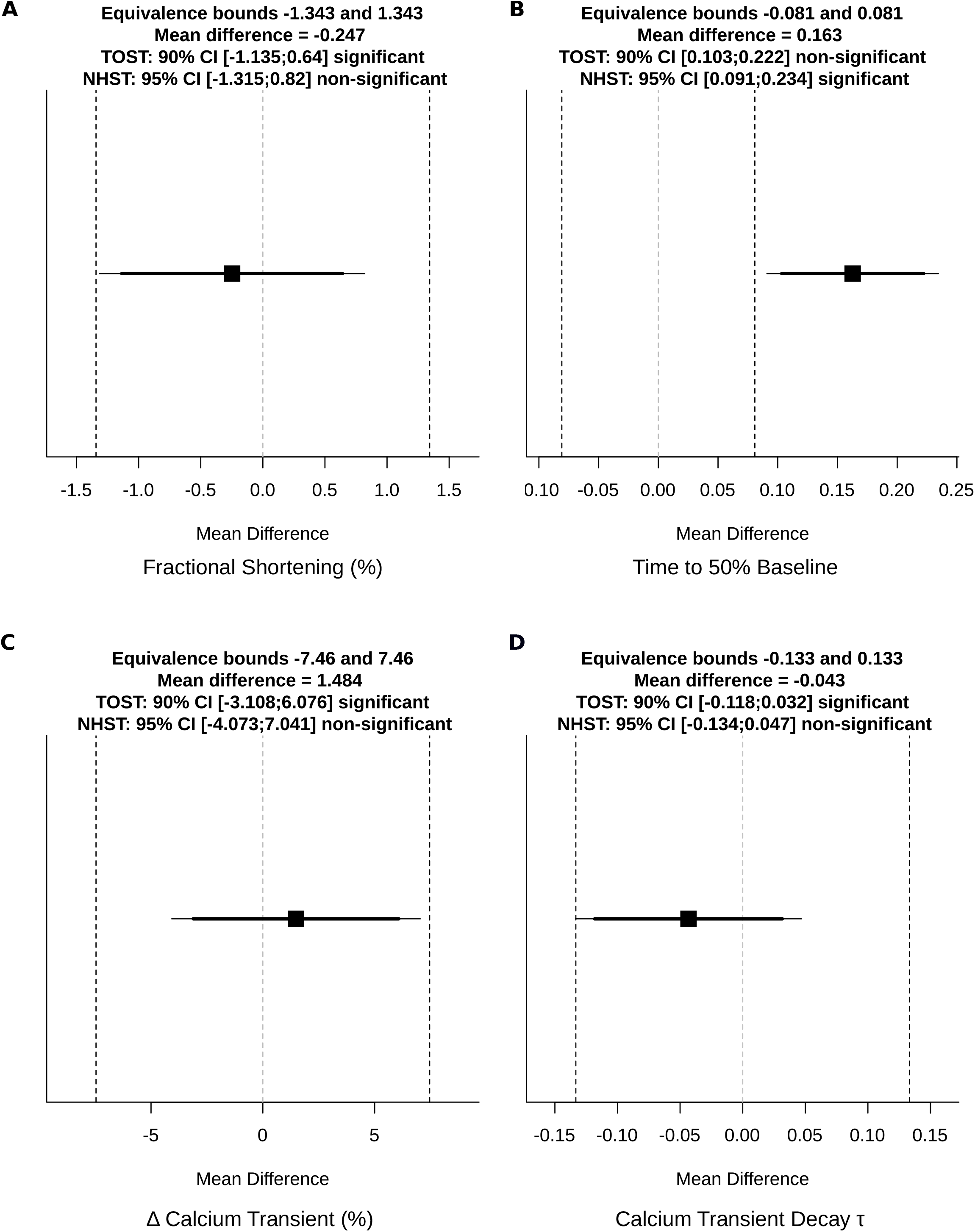
Raw test values for TOST and NHST tests in Figure 5. All figures are outputs of the TOSTER package in R. Equivalence bounds and mean differences are shown as raw bounds (i.e. in the original units of comparison).

