## Supplementary Methods for "Large particle fluorescence-activated cell sorting enables high quality single cell RNA-sequencing and functional analysis of adult cardiomyocytes"

**SUPPLEMENTARY MATERIALS**

***Mice***

For most experiments, we used adult (>3 months of age) C57BL/6J mice from Jackson Laboratory (Stock no. 000664). For mosaic experiments, we used the Ai9(RCL-tdT), obtained from Jackson Laboratory (Stock no. 007909). In this mouse, a loxP-flanked STOP casette prevents transcription of CAG promoter-driven tdTomato; this construct is inserted into the *Gt(ROSA)26Sor* locus. These mice are congenic to a C57BL/6J background. We refer to this mouse in the manuscript as Ai9. All animals were maintained compliant to protocols by the Johns Hopkins Animal Care and Use Committee.

***Langendorff Isolation of Cardiomyocytes***

Langendorff perfusion was performed based on established literature protocols[1,2]. We prepared the following buffers:

- Perfusion buffer: 120 mM NaCl, 5.4 mM KCl, 1.2 mM $NaH_2PO_4$, 20 mM $NaHCO_3$, 5.5 mM glucose, 5 mM BDM, 5 mM Taurine, and 1 mM $MgCl_2$, adjusted to pH 7.4
- Digestion buffer: 40 mL Perfusion buffer plus 35.8 mg Collagenase Type II (Worthington CLS-2), 3 mg Protease (Sigma P5147)
- Tyrode's buffer: 140 mM NaCl, 5 mM KCl, 10 mM HEPES, 5.5 mM glucose, and 1 mM $MgCl_2$, adjusted to pH 7.4

We used a horizontal (i.e. non-hanging) Langendorff apparatus with a chamber filled with perfusion buffer. To perform isolation, we first performed isofluorane anaesthesia on non-heparinized mice. Mice were observed until clearly anaesthetized and unresponsive to toe pinch, and subsequently euthanized by cervical dislocation. The heart was then rapidly excised from the chest and cannulated to the Langendorff apparatus. We flowed digestion buffer for 9 minutes at 1.5 mL/min. Subsequently, the ventricle was excised and minced. We filtered isolated cells through a 100 µM screen to eliminate large tissue chunks, spun down at 800 RPM for 1 minute (Eppendorf centrifuge 5702), and resuspended cells in 10 mL Tyrode's buffer.

For experiments where calcium-competent CMs were required, CMs were first resuspended in 10 mL Tyrode's with 1.5 µl of 1M $CaCl_2$. After 10 minutes, the CMs were resuspended in Tyrode's plus 2.5 µl of 1M $CaCl_2$. Subsequently, every 10 minutes, another 2.5 µl of 1M $CaCl_2$ until a total of 10 µL was added, resulting in a final concentration of 1 mM $Ca^{2+}$.

***Fluorescent Activated Cell Sorting***

We utilized a COPAS SELECT instrument (Union Biometrica) for the experiments in this manuscript. The COPAS SELECT was updated and rebranded as the FP-500, but the protocol used in the present study does not use the new features and thus the two are functionally indistinguishable. We optimized sorting for cardiomyocytes by using a sort delay of 8 and sort width of 6. Additionally, we used the following fluorescence settings: ext gain 50, green gain 200, yellow gain 200, red gain 255, extension integral gain 50, green integral gain 200, yellow integral gain 200, red integral gain 255, green PMT 800, yellow PMT 800, red PMT 1100. Coincidence check was selected to ensure proper single event sorting. We typically flowed cells between 20 – 60 events/second. All flow plots were saved as .lmd files and analyzed further using FlowJo v10. We maintained cells in Tyrode's buffer during the sort and sorted them into Tyrode's buffer; for calcium competent cells, we used Tyrode's + 1 mM $Ca^{2+}$. To run the machine, we used ClearSort Sheath Fluid (Sony, Lot 1218L345).

As an example of a commercial cell sorter, we tested the Sony SH800S cell sorter. We tested both 100 and 130 um microfluidic sorting chips. In a first study, we found that all objects that were sorted when using the 100 um chip were sheared; additionally, we encountered frequent clogs, as previously reported[3]. Thus, we subsequently focused experiments on the 130 um chip. All images presented in the manuscript were isolated after sorting through the 130 um chip.

### *Generation of tdTomato Mosaic Mouse*

To generate a tdTomato mosaic heart, we injected AAV9-cTNT-EGFP-T2A-iCre-WPRE vector (Vector Biolabs VB5413). For experiments, we used Ai9 Rosa-Tom heterozygotes at age P1. Prior to injection, we briefly anaesthetized pups on a crushed bed of ice for approximately 30 seconds. We then performed subcutaneous injection of virus ($4x10^{10}$ GC dose delivered in 30 µL). Mice were warmed for 5 minutes before being returned to the cage. The injected mice were observed regularly after the procedure to monitor health and recovery.

### *RNA Isolation for Quality Check*

To isolate RNA for quality check, we directly sorted CMs into Buffer RLT (Qiagen). Because each cell is sorted with approximately 0.5 – 1 µl volume, we sorted into a relatively large volume of RLT (approximately 3 – 5 mL, depending on number of sorted cells). RNA was immediately placed on dry ice and isolated as soon as possible. To isolate, we used the Rneasy Isolation Kit (Qiagen). Purified RNA was analyzed on the Advanced Analytical Fragment Analyzer.

### *scRNA-seq Library Preparation*

We performed scRNA-seq using the established SCRB-seq protocol[4]. Cells were sorted into 96-well plates composed of RNAse-free water plus 1:500 Phusion HF Buffer (New England Biolabs) and 1:250 RNAseOUT RNAse inhibitor(Life Technologies). Plates were placed on dry ice immediately after sorting and stored at -80C until ready for sequencing. For preparing sequencing libraries, we performed proteinase K treatment followed by RNA desiccation to reduce the reaction volume (50C for 15 minutes with plate seal on, followed by 95C for 10 minutes with plate seal removed). RNA was subsequently reverse transcribed using a custom template-switching primer as well as a barcoded adapter primer (see below). The customized SCRB-seq barcode primers contain a unique 6 base pair cell-specific barcode as well as a 10 base pair unique molecular identifier (UMI). Transcribed products were pooled and concentrated, with unincorporated barcode primers subsequently digested using Exonuclease I treatment (37C for 30 minutes, followed by 80C for 20 minutes to inactivate the enzyme). cDNA was PCR-amplified using Terra PCR Direct Polymerase (Takara Bio), performing 17 cycles of amplification. Final libraries were prepared using 1ng of cDNA per library with the Nextera XT kit (Ilumina) using a custom P5 primer as previously described. Pooled libraries were sequenced on two high-output lanes of the Illumina NextSeq500 with 16 base pair barcode read and 8 base pair i7 index read, and 66-92 base pair cDNA read design. The live/fixed study, which was performed first, utilized a 66 base pair read, which was sufficient for gene expression analysis and comparisons. However, we observed that with shorter read lengths, we obtained large counts of pseudogenes, many of which had significant sequence homology with mitochondrial genes. This likely implied mismapping due to short read length. Thus, for further studies (including the multi-chamber study presenting in the manuscript), we used the longer read length of 92 base pairs, which we found completely eliminated mapping to pseudogenes.

For the protocol, we used the following primers:

- Template-switching oligo: 5’-iCiGiCACACTCTTTCCCTACACGACGCrGrGrG-3’
- Barcoding primer: 5'-/5Biosg/ACACTCTTTCCCTACACGACGCTCTTCCGATCT[BC6]NNNNNNNNNNTTTTTTTTTTTTTTTTTTTTTTTTTTTTTTVN-3'
- PCR Amplification Primer: 5’-/5Biosg/ACACTCTTTCCCTACACGACGC-3’
- Custom i5 Oligo: 5’AATGATACGGCGACCACCGAGATCTACACTCTTTCCCTACACGACGCTCTTCCG*A*T*C*T*-3’ (* = phosphorothioate bond)

Additional details for the experiments in the manuscript are as follows:

- For the multi-chamber experiment – we initially sequenced 96 each of LV and RV CMs, and 72 each of LA and RA CMs. However, we found that the number of reads coming from the atrial samples was low. It was unclear whether this was due to poor atrial isolation or because atrial myocyte RNA content is lower. Regardless, we discarded the atrial data and focused on ventricular CMs.
- For the live/fixed experiment, we sequenced 36 each of live and fixed CMs.

***RNA-seq Analysis***

For the RNA-seq data generated in this manuscript, we mapped raw reads to the mouse genome GRCm38 from Ensembl concatenated with ERCC spike-in references. We mapped reads using zUMIs 2.2.3[5] with default settings and barcodes provided as a list. By default, zUMIs uses STAR (2.5.4b)[6] in a two-pass mapping format, and featureCounts through Rsubread (1.28.1) to tabulate counts and UMI tables. For the Delaughter and Nomura studies analysed in the manuscript, the raw fastq files were already demultiplexed by cell barcode. Thus, rather than use zUMIs, we mapped directly using STAR, using the same two-pass mapping and all other mapping settings as run by default through zUMIs. We subsequently counted using featureCounts, again maintaining settings as done during the zUMIs run. For the Gladka study, we were kindly provided with counts tables from the authors but could not acquire raw fastq files. Per the original manuscript, these tables were generated by mapping to the same mouse genome but using BWA-ALN; this distinction should be into account with regards to cross-study comparisons. For all studies, gene names were identified using biomaRt[7,8].

The data at each stage (fastq, counts, final analysis) is available on request, as is the code and R workspace to generate each of the figures. Briefly, each computational figure was generated as follows:

- Figure 3B – Using the multi-chamber dataset, we identified 126 cells that had achieved at least 200,000 reads but had fewer than 6000 genes at 200,000K reads (potential doublets) as a quality control cutoff. We then used SingleCellNet (0.1.0)[9]on the untransformed UMI counts. We used the Fluidigm C1 version of the Tabula Muris reference[10], filtering out only cell types that were annotated as being present in the heart to enable more specificity in analysis. We eliminated the “rand” category. The final plot was made using SingleCellNet’s built-in heatmap function.
- Figure 3C – We used the same 126 cells for the multi-chamber study as discussed above. For the Delaughter study, we initially analyzed cells from all timepoints. We used Seurat (2.4)[11] to cluster cells; doing so, we were able to clearly identify a population of CMs based on expression of markers such as *Myl2, Myh6, Myl3,* and *Tnni3*. We identified and focused in on

66 CMs at p21. We did not do any specific filtering for the Nomura dataset as the cells were previously identified under the microscope as CMs. For the Gladka data, we again used Seurat to cluster and identify a population of CMs in similar manner to with the Seidman data. We subsequently filtered out all CMs with fewer than 200,000 reads. Reads data as opposed to UMI data was used for both multi-chamber study and Gladka data set to enable true comparisons. For the Chevalier data, we eliminated MDX samples 4 and 5 as we found them to be extreme outliers regarding mitochondrial counts compared to other samples.

- Figure S2A – Candidate markers were selected for major cell types of interest. Plot was made using the pheatmap package in R, with column normalization to account for differences in read depth.
- Figure S2B – The same filtration criteria were used as in Figure 3C. We downsampled to approximately 200,000 reads using a custom R script and then computed the number of genes with any expression in each sample.
- Figures 3B/3C – Downsampled values were computed while mapping in zUMIs.
- Figure 3D – From Figures 3B and 3C, it is clear that there are differences between live and fixed cells both in gene detection and UMI detection at very high read depths. Our question was to determine whether, at these read depths, gene expression was at least well-correlated between live and fixed cells for detected genes. We selected 1.5M reads as our threshold of interest, and downsampled to 1.M reads from zUMIs (leaving behind 27 cells). We then averaged UMI expression both for the live and the fixed cells and plotted. To ensure that we were comparing genes that were genuinely expressed, we set a cutoff of 1 UMI to the means, and then plotted genes expressed both in live and fixed. The plot was made using base R plotting, and the $R^2$ value was computed by setting a linear model in R.

### *Ethanol Fixation*

To perform ethanol fixation of CMs, we first pelleted isolated CMs and resuspended in 200 µL of cold phosphate buffered saline (PBS). We then added 800 µL of -20C 100% ethanol drop by drop with gentle stirring to prevent clumping of cells. We performed experiments shortly after fixation, though we found that we could store cells up to two weeks at -20C without observing significant degradation of cell structure.

### *Function*

To perform functional analysis of CMs, we sorted CMs onto 7x10 mm laminin coated coverslips. Coverslips were then transferred to an inverted microscope (Nikon Eclipse TE-2000U). To measure sarcomere shortening and Ca2+ transient changes, cells loaded with the leak-resistant $Ca^{2+}$ indicator Fura-2AM (Molecular Probes) were studied using the IonOptix imaging system and software (IonOptix). Cells were paced at 1 Hz, and experiments performed at room temperature in Tyrode's containing 1 mM Ca2+ and 0.01% DMSO. To test for equivalence for functional assays, we used the R package TOSTER[12].

### *ImageJ Analysis*

We performed image analysis, particularly for Figures 1E and 1F, using Fiji through ImageJ2[13,14]. For assessing rod-shaped CMs, we used particle analysis. We found that the roundness metric effectively captured rod-shaped vs rounded/dead CMs, with a roundness of 0.43 (using 4x images) found empirically to separate rods from other cells.
